# Vasoactive Intestinal Peptide Amphiphile Micelle Material Properties Influence Their Cell Association and Internalization

**DOI:** 10.1101/2024.06.10.598391

**Authors:** Xiaofei Wang, Agustin T. Barcellona, Fateme Nowruzi, Kambrie Brandt, Megan C. Schulte, Luke E. Kruse, Eric Dong, Bret D. Ulery

## Abstract

Vasoactive intestinal peptide (VIP) is a promising anti-inflammatory peptide therapeutic that is known to induce biological effects by interacting with its cognate receptor (*i.e.*, VPAC) on the surface of antigen presenting cells (APCs). For VIP-based drug delivery technologies like nano- and microparticles, little is known about the effect VPAC targeting has on APC behavior. This is further influenced by the fact that particulate material properties including chemistry, shape, and size are all known to influence APC behavior. In this study, peptide amphiphile micelles (PAMs) were employed as a modifiable platform to study the impact VPAC targeting and physical particle properties have on their association with macrophages. VIP amphiphile micelles (VIPAMs) and their scrambled peptide amphiphile micelle analogs (^S^VIPAMs) were fabricated from various chemistries yielding particle batches that were comprised of spheres (10 - 20 nm in diameter) and/or cylinders of varying lengths (*i.e.*, 20 - 9000 nm). Micelle surface attachment to and internalization by macrophages were observed using confocal microscopy and their association was characterized by flow cytometry. The enclosed work provides strong evidence that macrophages rapidly bind VPAC specific micelles independent of physical properties though micelle shape and size as well as receptor-specificity all influence their long-term macrophage association. Specifically, a mixture of spherical and short cylindrical VIPAMs were able to achieve the greatest cell association which may correlate to their capacity to fully bind the VPAC receptors available on the surface of macrophages. These results provide the foundation of how nano- and microparticle physical properties and targeting capacity synergistically influence their capacity to associate with APCs.

## Introduction

Vasoactive intestinal peptide (VIP) is an endogenous neuropeptide that has emerged as a promising immunomodulatory candidate for treating a wide variety of inflammatory conditions^1^. VIP adeptly mitigates proinflammatory cytokine expression, promotes anti-inflammatory activity, and modulates immune cell processes, contributing to a balanced immune homeostasis^2,3^. Despite its safety and efficacy, the clinical development of VIP as an immunotherapeutic agent faces several challenges including possessing a very short half-life and undesirable off-target toxicity^4^. To overcome these limitations, researchers have begun engineering VIP using various approaches to improve its pharmaceutical properties while maintaining or enhancing its immunomodulatory activity^5^. In the context of peptide therapeutics, lipidation has been found to enhance peptide delivery and bioactivity^6,7^ making this a promising approach for VIP delivery. Despite the successes of lipid-peptide drugs, further investigation into the structure and properties of these is crucial to understanding their effects and biofunctionality.

Peptide amphiphiles (PAs) are lipid-modified peptides that self-assemble into nanostructured peptide amphiphile micelles (PAMs) in water. PAMs offer several advantages that overcome the limitations associated with free peptide administration; they can prevent peptidyl diffusion^8^, increase local peptide concentration^9^, enhance intracellular peptide delivery^10^, and improve peptide bioactivity^11^. The effects of PAMs on the immune system depend on different factors, including, but not limited to, the shape and size of micelles^12,13^. Specifically, these factors can affect micelle-cell association and internalization, contributing to the activation of antigen presenting cells (APCs) to an inflammatory state^14,15^. Therefore, the investigation of macrophage association with and uptake of micelles of different shapes and sizes is crucial to understanding how they affect the immune system. Moreover, most studies have focused on the cell processing of solid particles (*i.e.*, polymeric^16,17^, silica^18,19^, and metallic^20,21^) which may behave differently than supramolecular particles like PAMs^22^. Importantly, few studies analyzing VIP amphiphile micelle (VIPAM) structure and shape as well as their effects on the immune system and cell interactions have been undertaken. Recently, our research group synthesized a small library of VIPAMs containing different N-terminal palmitoyl and dipalmitoyllysine moieties and zwitterion-like lysine - glutamic acid peptide segments which includes PalmK-(EK)_4_-VIP, PalmK-VIP-(KE)_4_, Palm_2_K-(EK)_4_-VIP, and Palm_2_K-VIP-(KE)_4_. Morphological assessments demonstrated that the zwitterion-like block location within the PAs greatly impacted the assembly, structure, and charge of VIPAMs which influenced their stability, cell interactions, and bioactivity^23^. Based on this work, external zwitterion-like block-VIPAs were selected for further study due to their optimized shape, size, charge, and durable anti-inflammatory effects.

This study further investigates the intricate interplay between the physical properties of VIPAMs and their cellular interactions with macrophages. The strategic design of PAMs with external zwitterion-like lysine - glutamic acid sequences and dipalmitoyllysine additions was employed to help optimize VIP delivery and bioactivity. Specifically, structurally unique VIPAMs were formed with shapes ranging from small spheres to long cylinders. VIPAM shapes were controlled by altering both the ratio of lysine to glutamic acid within the (KE)_x_ sequences as well as the number of hydrocarbon tails in the lipid moiety. Interestingly, micelle shape and size directly influenced cellular association and internalization. This effect demonstrated the clear benefit of utilizing PAMs to effectively deliver VIP to target cells for immunomodulation. Additionally, these findings open new possibilities for developing advanced immunotherapies leveraging the unique functional capacity of PAMs.

## Materials and Methods

Vasoactive intestinal peptide (VIP, Ac**-**HSDAVFTDNYTRLRKQMAVKKYLNSILN-CONH_2_), scrambled VIP (^S^VIP, Ac-NSDLIATDSYTRMRKQVLANKKFHYLVN-CONH_2_), Fmoc-VIP-KEKEKEKE-CONH-Resin, Fmoc-^S^VIP-KEKEKEKE-CONH-Resin, Fmoc-VIP-KEKEKKKK-CONH-Resin, Fmoc-^S^VIP-KEKEKKKK-CONH-Resin, Fmoc-VIP-KEKEEEEE-CONH-Resin, and Fmoc-^S^VIP-KEKEEEEE-CONH-Resin were purchased from Synpeptide Co., Ltd, China. Fmoc-Lys(Fmoc)-OH, Fmoc-Lys(ivDde)-OH, and ivDde-Lys(Fmoc)-OH were acquired from Novabiochem. AK Scientific, Inc. and AnaSpec, Inc. were the suppliers used for 2- (1H-benzotriazol-1-yl)-1,1,3,3-tetramethyluronium hexafluorophosphate (HBTU) and 1-hydroxybenzotriazole hydrate (HOBt hydrate), respectively. Palmitic acid (Palm) and 5(6)- carboxyfluorescein (FAM) were acquired from Acros Organics. Piperidine, triisopropylsilane (TIS), N,N-diisopropylethylamine (DIEA) were purchased from Chem-Impex International, Inc,. Dimethylformamide (DMF), 1-methyl-2-pyrrolidinone (NMP), acetic anhydride (Ac_2_O), hydrazine monohydrate, thioanisole (TA), trifluoroacetic acid (TFA), phenol, ethanedithiol (EDT), and diethyl ether were bought from Millipore Sigma. A glass peptide reaction vessel was obtained from Chemglass Life Sciences. Fetal bovine serum (FBS) was sourced from Sigma Aldrich, located in St. Louis, MO. Penicillin/streptomycin solution was purchased from ThermoFisher, based in Waltham, MA. Lipopolysaccharide (LPS) was obtained from Santa Cruz Biotechnology, Inc., Dallas, TX. The molecule 1,6-diphenyl-1,3,5-hexatriene (DPH) was acquired from Sigma Aldrich, St. Louis, MO.

### Solid Phase Peptide Synthesis (SPPS) and Lipidation

The library of VIP and ^S^VIP products were synthesized on a resin support employing standard Fmoc solid phase peptide synthesis (SPPS) using our previously reported method^23^. The general approach for the synthesis of the PA formulations is outlined in **Scheme 1**. The dry resin was rinsed with NMP for 2 hours under a nitrogen atmosphere with bubbling to provide mixing. Between each step, the resin was pre-washed three times with the solvent for the next coupling or deprotection reaction. Fmoc deprotection was achieved by treatment with 25% piperidine in DMF (2 x 30 min). The ivDde protecting group was removed using 2% hydrazine monohydrate in DMF (6 x 20 min). For amino acid or lipid conjugation, Fmoc-Lys(ivDde)-OH, ivDde-Lys(Fmoc)-OH, Fmoc-Lys(Fmoc)-OH, or palmitic acid (1 equiv) were pre-activated with HBTU (4.2 equiv), HOBt (5 equiv), and DIEA (10 equiv) in NMP for 10 min. The activated species were then coupled to the N-terminus of the resin-bound peptide over three 90 min coupling cycles. For fluorescent labeling, FAM (1 equiv) was pre-activated similarly using HBTU/HOBt/DIEA and coupled to the N-terminus over three overnight cycles. Remaining unreacted N-termini were capped by treatment with 5% acetic anhydride and 7% DIEA in NMP for 15 min.

**Scheme 1.**
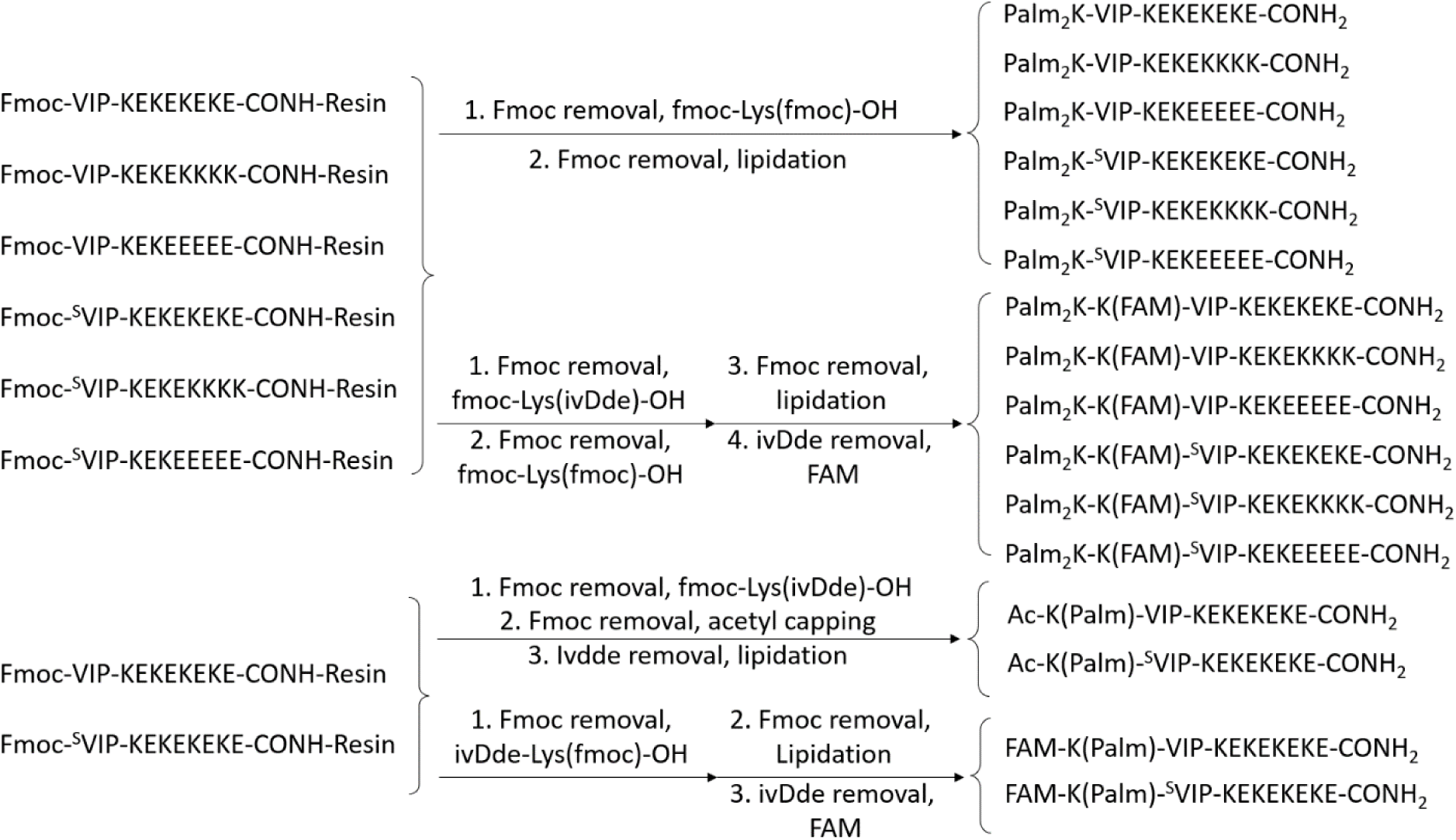
Fluorophore labeled and non-fluorophore labeled VIP amphiphiles and ^S^VIP amphiphiles synthesis approaches.

After modifications, the resin was washed with methanol (3x), transferred to a 15 mL centrifuge tube as a methanol slurry, and dried under high vacuum. Peptide cleavage from the resin and global deprotection of side chains was achieved by a 2 h treatment with a cleavage cocktail of TFA/triisopropylsilane/water/phenol/ethanedithiol (87.5:2.5:2.5:2.5:2.5:2.5). Cleaved peptides were precipitated in diethyl ether and then centrifuged. The solid was collected, resuspended in fresh diethyl ether by vortexing (3x) to wash the solids, and finally dissolved in minimal deionized water for lyophilization to yield the purified product as a powder. All crude VIP and VIP amphiphiles (VIPAs) as well as their scrambled analogues were analyzed by an analytical high-pressure liquid chromatograph (HPLC, Beckman Coulter) and purified by preparative mass-spectrometry fraction-controlled HPLC (HPLC-MS) on a C4 column (Milford, MA).

### Critical Micelle Concentration (CMC)

A critical micelle concentration (CMC) assay was performed to determine the concentration at which different VIPAs and ^S^VIPAs form micelles in. In this study, DPH was employed as a fluorophore to determine the CMC. DPH can exhibit weak fluorescence in aqueous environments, but becomes strongly fluorescent when incorporated within a hydrophobic domain. PAs were serially diluted (31.6 μM to 0 μM) in 1 µM DPH in PBS and allowed to incubate for at least 1 hour. Sample fluorescence was measured by a Cytation 5 fluorospectrophotometer (BioTek Instruments, Inc., Winooski, VT) at *ex*: 350 nm, *em*: 428 nm. The jerk point between stable low fluorescence and the beginning of a rapid increase in fluorescence was considered the CMC for the sample.

### Micelle Morphology Characterization

PAM morphology was characterized by negative-stain transmission electron microscopy (TEM) using a previously established method^23^. Micelle solution (5 µM) was added to a carbon support TEM grid (200 mesh, Electron Microscopy Sciences, Hatfield, PA). After 5 minutes of incubation, the solution was removed and immediately followed by the addition of 5 µL of NanoW (Nanoprobes, Inc, Yaphank, NY). After 3 minutes of incubation, the solution was removed and grids were left to dry followed by imaging with a JEOL JEM-1400 TEM at 120 kV. Images of at least three different spots on each grid were taken and analyzed.

### Secondary Structure Measurement

Micelle secondary structure was assessed by circular dichroism (CD) using a Chirascan V100 CD spectrometer from Applied PhotoPhysics (Leatherhead, UK). Micelle solutions (40 μM) were prepared in PBS and loaded into a 0.5 mm cuvette and measured spectrophotometrically from 200 nm to 250 nm with an interval of 1 nm. The data were fit using a linear combination of polylysine and polyglutamine structures to calculate approximate α-helical, β-sheet, and random coil content.

### Cell Assessment

Immunomodulatory experiments were conducted with murine RAW 264.7 (macrophage-like cells - Mφs) using an established protocol^23^. Complete Dulbecco’s Modified Eagle Medium (DMEM) was made by supplementing DMEM with 1% penicillin/streptomycin (ThermoFisher, Waltham, MA), and 10% FBS (FBS, Sigma Aldrich, St. Louis, MO). Mφs were cultured in 100 mm diameter, non-treated petri-dishes in complete DMEM at 37 ℃ and 5% CO_2_. The cells were seeded in non-treated 24-well plates at 50,000 cells per well. After overnight incubation in complete DMEM, cells were stimulated with 0.1 µg/mL of LPS (Santa Cruz Biotechnology, Inc., Dallas, TX) along with varying concentrations of VIP, VIPAs, ^S^VIP, or ^S^VIPAs. All peptides were non-fluorophore labeled whereas micelle products possessed small amounts of fluorophore labeled components (*i.e.*, 1:19 FAM-PA:Non-FAM-PA). Cells given media alone without the LPS stimulus were studied as a negative control. After 1 or 6 hours of incubation, supernatant solutions were collected for assessment of secreted tumor necrosis factor-α (TNF-α) content using an enzyme-linked immune sorbent assay (ELISA) kit (Biolegend, San Diego, CA). Cells were washed with PBS to delaminate them from the bottom of the non-treated 24-well plates after which a second wash with PBS was performed to ensure the collection of all cells. To analyze viability, the collected cells were stained with the Live-or-Dye 594/614 Fixable Viability Staining Kit^TM^ (Biotium, Fremont, CA) and then analyzed by a flow cytometer (BD LSRFortessa X20) equipped with FACSDiva 8.0 Software to detect the fluorescence intensity of each cell. FlowJo was utilized to gate the cells and analyze the fluorescence signals.

For microscopy studies, cover slips were added to the non-treated 24-well plates before 0.05 million cells were placed in each well. After overnight incubation, cells were treated with 0.1 µg/mL of LPS containing the same VIPAs or ^S^VIPAs possessing small amounts of fluorophore labeled components (*i.e.*, 1:19 FAM-PA:Non-FAM-PA) as was used for the previous studies. After 1 hour or 6 hours of incubation, cells bound to coverslips were washed with Hank’s Balanced Salt Solution (HBSS) and stained with 5 µg/mL wheat germ agglutinin (WGA) for 10 minutes at 37 ℃ and 5% CO_2_. Cells were then washed twice with PBS to remove any non-associated stain. Cells were fixed through exposure to 4% paraformaldehyde (PFA) for 15 minutes. A total of three PBS washes were used to get rid of any excess PFA and then cells were subjected to 1% Triton X-100 for 15 minutes to help permeabilize the fixed cell membrane. After washing the cells three times with PBS, they were preincubated with 1% bovine serum albumin (BSA) in PBS for 30 minutes to prevent non-specific binding of the 4,6’-diamidino-2-phenylindole (DAPI) stain used to identify the cell nucleus. DAPI-containing mounting solution was then used to adhere the cover slip to glass slides that were then analyzed for their fluorescence using a Leica TCS SP8 STED and MP confocal microscope. Leica LAS X 3D Analysis software was employed to analyze, compile, and present the fluorescent data from the three channels used (*i.e.*, 420 - 480 nm, 500 - 550 nm, and 670 nm - 730nm).

### Statistical Analysis

The analysis of group comparisons was conducted using JMP software from SAS Institute (Cary, NC). This involved performing an analysis of variance (ANOVA) assessment followed by Tukey’s honest significant differences (HSD) test to identify pairwise statistically significant differences (p < 0.05). In visual representations, groups with distinct letters or marked by a * signify statistically significant differences in means, whereas groups with the same letter or no symbol suggest statistically insignificant differences.

## Results and Discussion

### Toxicity and Bioactivity Screen of VIP and VIPAMs

In order to investigate the impact micelle structural changes have on Mφ association, the previously studied double lipid, external zwitterion-like peptide block VIPA formulation (*i.e.*, Palm_2_K-VIP-(KE)_4_) was selected as a starting PA to design new products from as it displayed interesting concentration-, shape-, and size-dependent TNF-α expression modulation behavior, especially when compared to its long cylindrical, single lipid analog (*i.e.*, PalmK-VIP-(KE)_4_)^23^. An initial dose effect study was undertaken to find the optimal concentration for these two VIPAMs as well as their unlipidated, unmicellized peptide controls above the lower limit of bioactivity (*i.e.*, 1 µM), but below a potential cytotoxicity upper limit (*i.e.*, 10 µM). LPS-treated Mφs were exposed to 1, 2.5, 5, or 10 µM VIP, PalmK-VIP-(KE)_4_, or Palm_2_K-VIP-(KE)_4_ for six hours, and cell viability was subsequently assessed using flow cytometry (**Figure 1a**). Minimal cytotoxicity was induced by exposure of cells up to 5 µM VIP, but cells were found to be only 70 - 80 % as viable as untreated cells when incubated with 10 µM PalmK-VIP-(KE)_4_ or Palm_2_K-VIP- (KE)_4_. TNF-α expression of LPS-activated Mφs exposed to the same concentration gradient of VIP and VIPAMs was also measured (**Figure 1b**). No appreciable modulation of TNF-α secretion was seen with VIP or VIPAMs at the lowest two concentrations (*i.e.*, 1 µM and 2.5 µM), but significant TNF-α suppression was observed for VIP and Palm_2_K-VIP-(KE)_4_ at the highest two concentrations (*i.e.*, 5 µM and 10 µM). Based on these findings, an optimized VIPAM working concentration of 5 µM was chosen to explore the influence micelle physical properties had on their cell association and internalization.

**Figure 1.**
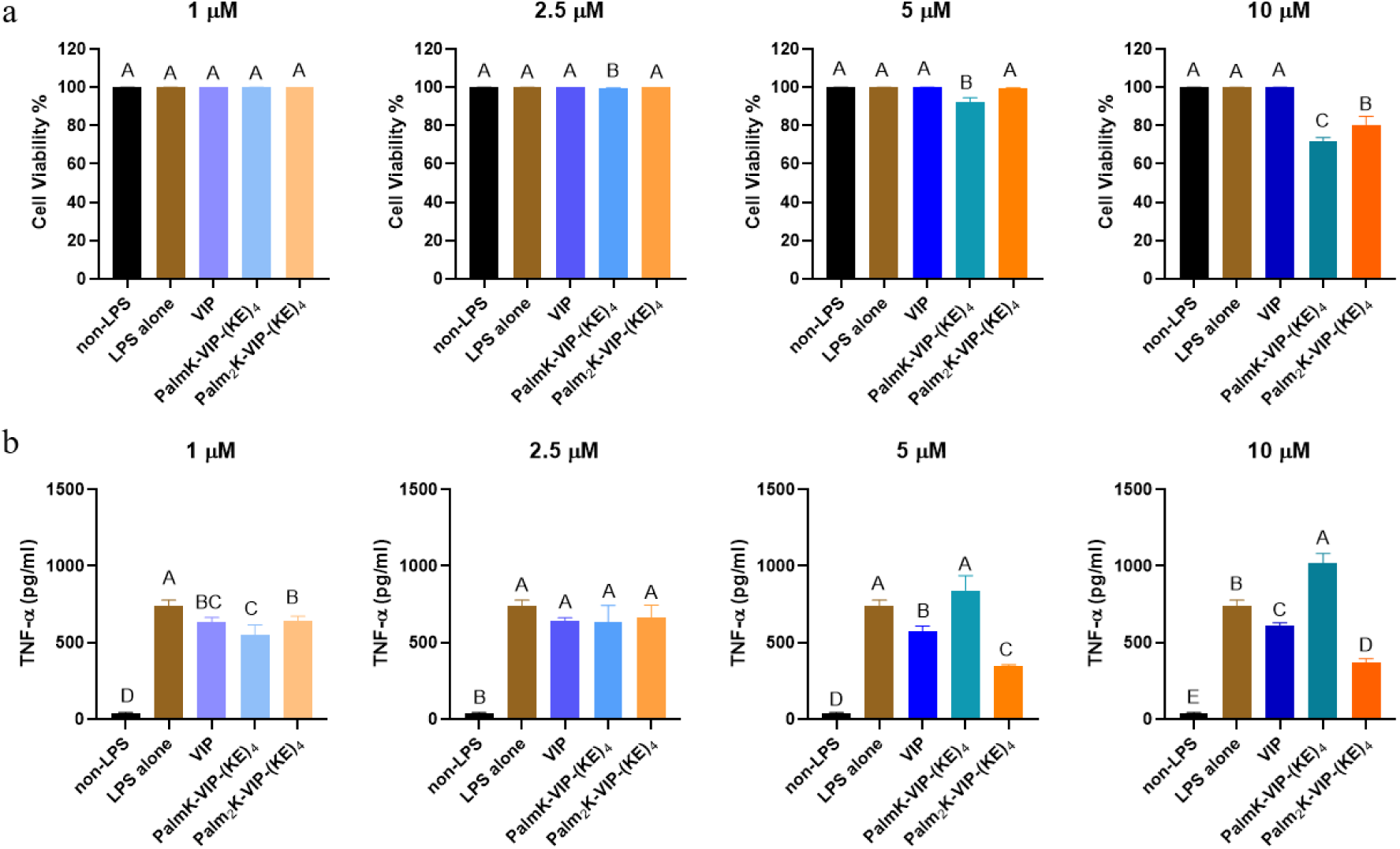
Effect of 6 h VIPAM exposure on LPS-activated Mφ cell health (a) and TNF-α secretion (b). In each graph, groups possessing the same letter have no statistically significant difference (p > 0.05).

### Chemical Structure and Physical Properties of VIPAs and ^S^VIPAs

Structurally distinct VIPAM formulations homologous to Palm_2_K-VIP-(KE)_4_ were then generated in order to study the influence micelle architecture has on Mφ association and uptake. Incorporation of several lysine residues has been shown to reduce micelle size^24^ likely due to corona-based charge repulsion increasing micelle surface curvature yielding spherical micelles in accordance with Israelachivili’s surfactant theory^25^. Specifically, this theory defines a critical packing parameter (CPP) of surfactant molecules like PAs as:

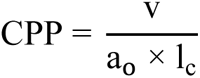

where v represents the hydrocarbon core volume, a_o_ is the effective head group area, and l_c_ is the hydrocarbon chain length. Based on this concept, the VIPAM hydrophilic peptide block (*i.e.*, (KE)_4_) was modified to contain short charge repulsive regions at the peptide C-terminus. Specifically, the last four amino acids were made all positive (*i.e.*, -(KE)_2_K_4_ – zwit-cat) or all negative (*i.e.*, -(KE)_2_E_4_ – zwit-an) yielding PAs with sequences of Palm_2_K-VIP-(KE)_2_K_4_ and Palm_2_K-VIP-(KE)_2_E_4_.

The CMCs of the four VIPAs of interest (*i.e.*, PalmK-VIP-(KE)_4_, Palm_2_K-VIP-(KE)_4_, Palm_2_K- VIP-(KE)_2_K_4_ and Palm_2_K-VIP-(KE)_2_E_4_) were obtained to determine the minimal PA concentrations necessary for micelle formation as well as to observe the relationship between PA chemical structure and micellization. (**Figure 2a-d**).

**Figure 2.**
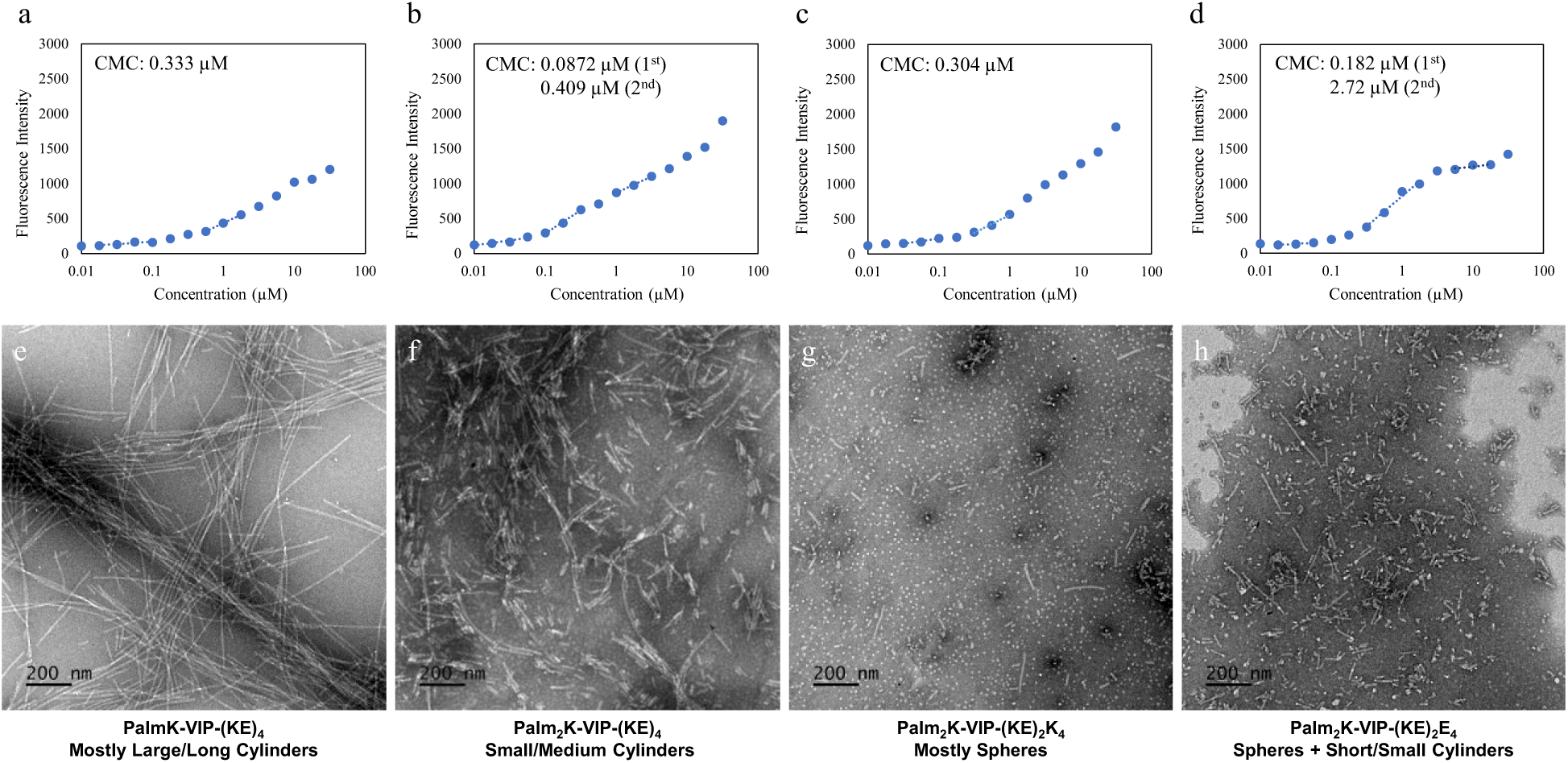
Critical micelle concentration (CMC) and micelle morphology of PalmK-VIP-(KE)_4_ (a, e), Palm_2_K-VIP-(KE)_4_ (b, f), Palm_2_K-VIP-(KE)_2_K_4_ (c, g) and Palm_2_K-VIP-(KE)_2_E_4_ (d, h). All micrographs were taken at a magnification of 12,000 x with the scale bar for all images set at 200 nm.

All four VIPAs possessed relatively low CMCs with two formulations (*i.e.,* Palm_2_K-VIP- (KE)_4_ and Palm_2_K-VIP-(KE)_2_E_4_) exhibiting a second CMC possibly indicating a secondary micelle structure transition. When looking at the initial CMC values for each formulation, single lipid VIPAs (*i.e.,* PalmK-VIP-(KE)_4_) had the largest first CMC (**Figure 2a** – 0.333 µM). Palm_2_K- VIP-(KE)_4_ had a lower CMC (**Figure 2b –** 0.0872 µM) than double lipid VIPAs possessing uneven ratios of either lysines or glutamic acids (*i.e.,* **Figure 2c -** Palm_2_K-VIP-(KE)_2_K_4_ - 0.304 µM and **Figure 2d** - Palm_2_K-VIP-(KE)_2_E_4_ - 0.182 µM)potentially arising from charge repulsion between the hydrophilic head groups^26^. Excitingly, all VIPA formulations formed micelles at the identified working concentration (*i.e.*, 5 µM), so the morphology of all VIPAMs at this concentration was assessed using TEM. To assist in describing nanoparticle shape, micelle architecture was divided into six categories: spheres (L/D ∼ 1), short cylinders (3 < L/D ≤ 10), small cylinders (10 < L/D ≤ 30), medium cylinders (30 < L/D ≤ 100), large cylinders (100 < L/D ≤ 300), and long cylinders (300 < L/D). It was found that PalmK-VIP-(KE)_4_, Palm_2_K-VIP-(KE)_4_, Palm_2_K-VIP-(KE)_2_K_4_ and Palm_2_K-VIP-(KE)_2_E_4_ formed mostly large/long cylinders (1000 - 9000 nm in length), small/medium cylinders (100 - 700 nm in length), mostly spheres (10 - 15 nm in length), and a mix of spheres (10 - 20 nm in diameter) and short/small cylinders (20 - 300 nm in length), respectively (**Figure 2e-h**). Single lipid VIPAs (*i.e.*, PalmK-VIP-(KE)_4_) assembled into more elongated structures (**Figure 2e**) compared to double lipid VIPAs (*i.e.*, Palm_2_K-VIP-(KE)_4_, Palm_2_K-VIP-(KE)_2_K_4_, and Palm_2_K-VIP-(KE)_2_E_4_) (**Figure 2f-h**). The greater steric hindrance found in the hydrophobic core of double lipid VIPAMs in contrast to the single lipid VIPAMs likely resulted in much looser amphiphile packing causing the effective head group area (a_o_) to be greater than the doubling of the hydrocarbon core volume (v) leading to overall lower CPP values. This observation is similar to previously published results on the influence lipid number has on micelle size^11^. Furthermore, zwit-cat double lipid VIPA (*i.e.*, Palm_2_K-VIP-(KE)_2_K_4_) and zwit-an double lipid VIPA (*i.e.*, Palm_2_K-VIP-(KE)_2_E_4_) assembled into more compact micelles (*i.e.*, **Figure ^2^g-h** - at least some population of spheres were found with both formulations) than zwit double lipid VIPA (*i.e.*, **Figure 2f** - Palm_2_K-VIP-(KE)_4_ - only cylindrical micelles). This result suggested that corona-based charge repulsion was likely able to further increase effective head group area (a_o_) leading to even greater micelle curvature.

In addition to the four VIPAM formulations, a scrambled VIP peptide (^S^VIP) was synthesized and utilized to generate respective ^S^VIPAM analogs to explore the influence peptide specificity has on micelle association and internalization by cells. In order to generate a non-functional, biocompatible scrambled peptide control, an algorithm employing a Boltzmann Factor scoring function was first utilized to rearrange the native VIP peptide sequence. Specifically, amino acids known to participate in receptor binding were rearranged with residues possessing similar hydropathy, while minimal modifications were performed for amino acids which contribute to overall VIP structural integrity^27^. The final sequence of ^S^VIP peptides and how they compare to VIP can be found in **Table 1**. Cytotoxicity and bioactivity studies were performed with ^S^VIP and VIP at 5 µM using LPS-treated Mφs along with LPS-treated cells with no stimulus and non-LPS treated cells (**Figure 3**). The viability of ^S^VIP-treated cells remained high, though their secreted TNF-α concentration was slightly higher than those exposed to LPS alone possibly due to formation of amorphous aggregates at 5 µM (**Figure 4**) which could be responsible for inducing mild, non-specific inflammation in Mφs^28,29^. In contrast, VIP lacks any particle structure when observed by TEM (*data not shown*). Upon formation of an appropriate ^S^VIP peptide, a series of amphiphiles with the same chemical composition as the aforementioned VIPAs were produced by replacing the VIP block with ^S^VIP producing non-specific, scrambled micelles (^S^VIPAMs) for further investigation (**Table 1**).

**Figure 3.**
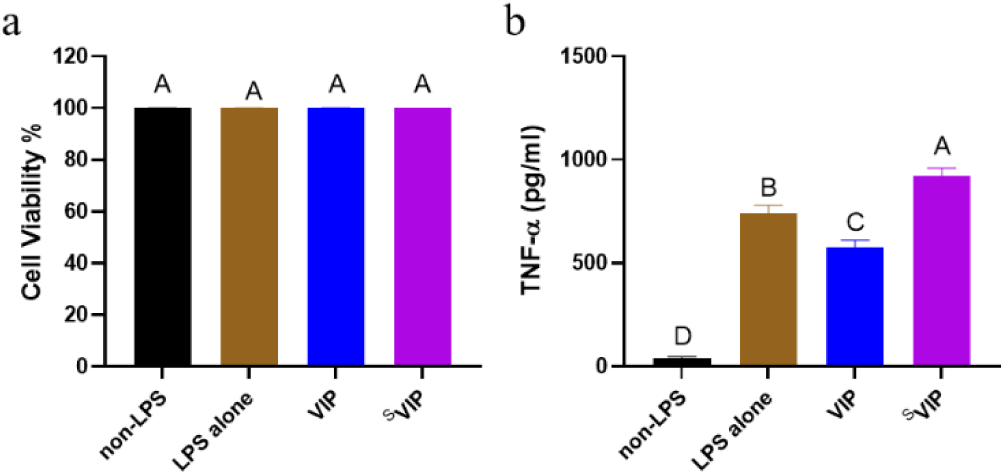
Influence of 6-hour 5 μM ^S^VIP exposure on LPS-activated Mφ cell health (a) and TNF-a secretion (b). In each graph, groups possessing the same letter have no statistically significant difference (p > 0.05).

**Figure 4.**
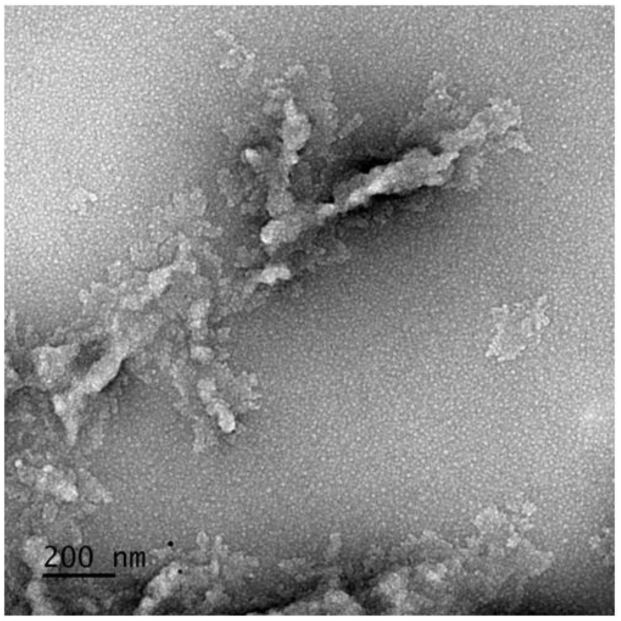
Amorphous aggregates formed with 5 µM ^S^VIP. The micrograph was taken at a magnification of 12,000 x with the scale bar representing 200 nm.

**Table 1.**
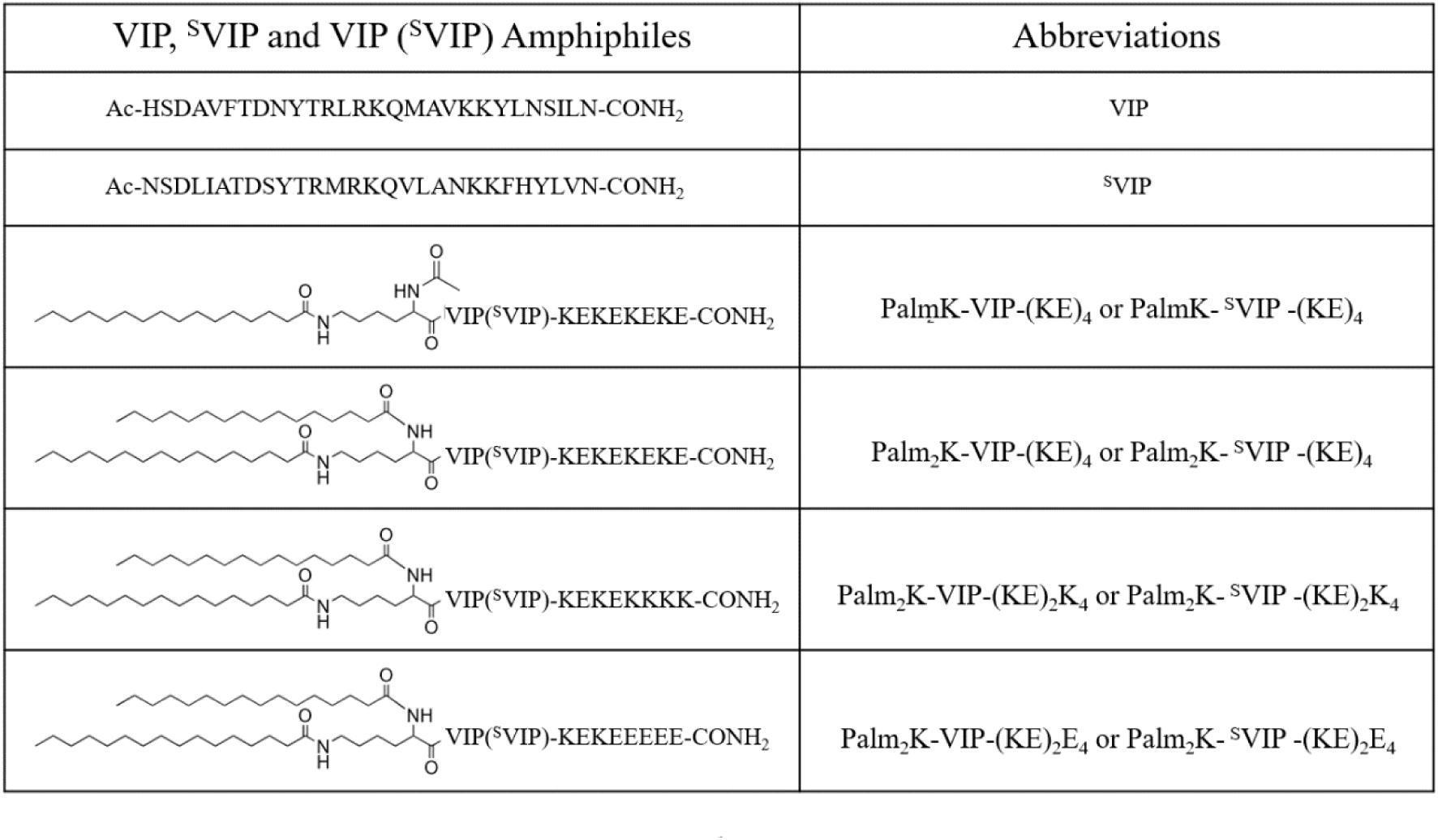
Sequences of VIP and ^S^VIP and chemical structures of VIPAs and ^S^VIPAs.

The CMCs and TEM images of the four ^S^VIPA analogs were determined to better understand the influence bioactive peptide chemical structure has on micelle formation and architecture (**Figure 5a-h**). The physical characterization results for all VIPA and ^S^VIPA formulations are summarized in **Table 2** to allow for easier comparisons among the data across different parameters. The micellization behavior of ^S^VIPA formulations was relatively similar to that of VIPAs. The CMC of the single lipid ^S^VIPA (*i.e.*, PalmK-^S^VIP-(KE)_4_, **Figure 5a -** 2.85 µM) was found to be higher than its double lipid ^S^VIPA analog (*i.e.*, Palm_2_K-^S^VIP-(KE)_4_, **Figure 5b** −0.158 µM) which follows the same convention found for VIPAs. Additionally, two CMCs were observed for zwit-an double lipid ^S^VIPA (*i.e.*, Palm_2_K-^S^VIP-(KE)_2_E_4_, **Figure 5d**) similar to its corresponding VIP formulation (*i.e.*, Palm_2_K-VIP-(KE)_2_E_4_, **Figure 2e**). Interestingly, zwit double lipid ^S^VIPA (*i.e.*, Palm_2_K- ^S^VIP-(KE)_4_, **Figure 5b**) and zwit-cat double lipid ^S^VIPA (*i.e.*, Palm_2_K- ^S^VIP-(KE)_2_K_4_, **Figure 5c**) exhibited one and two CMCs, respectively, which is inverted from what was found for their analogous VIP formulations (*i.e.*, **Figure 2b** - Palm_2_K- VIP-(KE)_4_ and **Figure 2d** - Palm_2_K- VIP-(KE)_2_K_4_). Regardless of these differences, all ^S^VIPA formulations were found to have formed micelles at the 5 µM working concentration like their VIPA counterparts. The similarities and differences detected in the CMC profiles between VIPAs and ^S^VIPAs with the same N-terminal and C-terminal chemical modifications was also reflected in their morphologies. TEM images of PalmK-^S^VIP-(KE)_4_, Palm_2_K-^S^VIP-(KE)_4_, Palm_2_K-^S^VIP-(KE)_2_K_4_, and Palm_2_K- ^S^VIP-(KE)_2_E_4_ revealed their micellar morphologies to be mostly large/long cylinders (1000 - 9000 nm in length), spheres (10 - 15 nm in diameter), small/medium cylinders (100 - 400 nm in length), and a mix of spheres (10 - 20 nm in diameter) and short/small cylinders (50 - 250 nm in length), respectively (**Figure 5e-h**). Single lipid ^S^VIPA (*i.e.*, **Figure 5e** - PalmK-^S^VIP-(KE)_4_) and zwit-an double lipid ^S^VIPA (*i.e.*, **Figure 5h** - Palm_2_K-^S^VIP-(KE)_2_E_4_) possessed similar architectures as their VIPA counterparts (**Figure 2e** and **Figure 2h**). Interestingly, the other two ^S^VIPA formulations flipped micelle shapes with their VIPA counterparts with compact micelles generated by zwit-cat double lipid ^S^VIPA (**Figure 5g**) and zwit double lipid VIPA (**Figure 2f**) whereas mostly spherical micelles were present with zwit double lipid ^S^VIPA (**Figure 5f**) and zwit-cat double lipid VIPA (**Figure 2g**). This inversion mimicked what was observed with CMCs further supporting the semi-stable spherical micelle shape found with some formulations.

**Figure 5.**
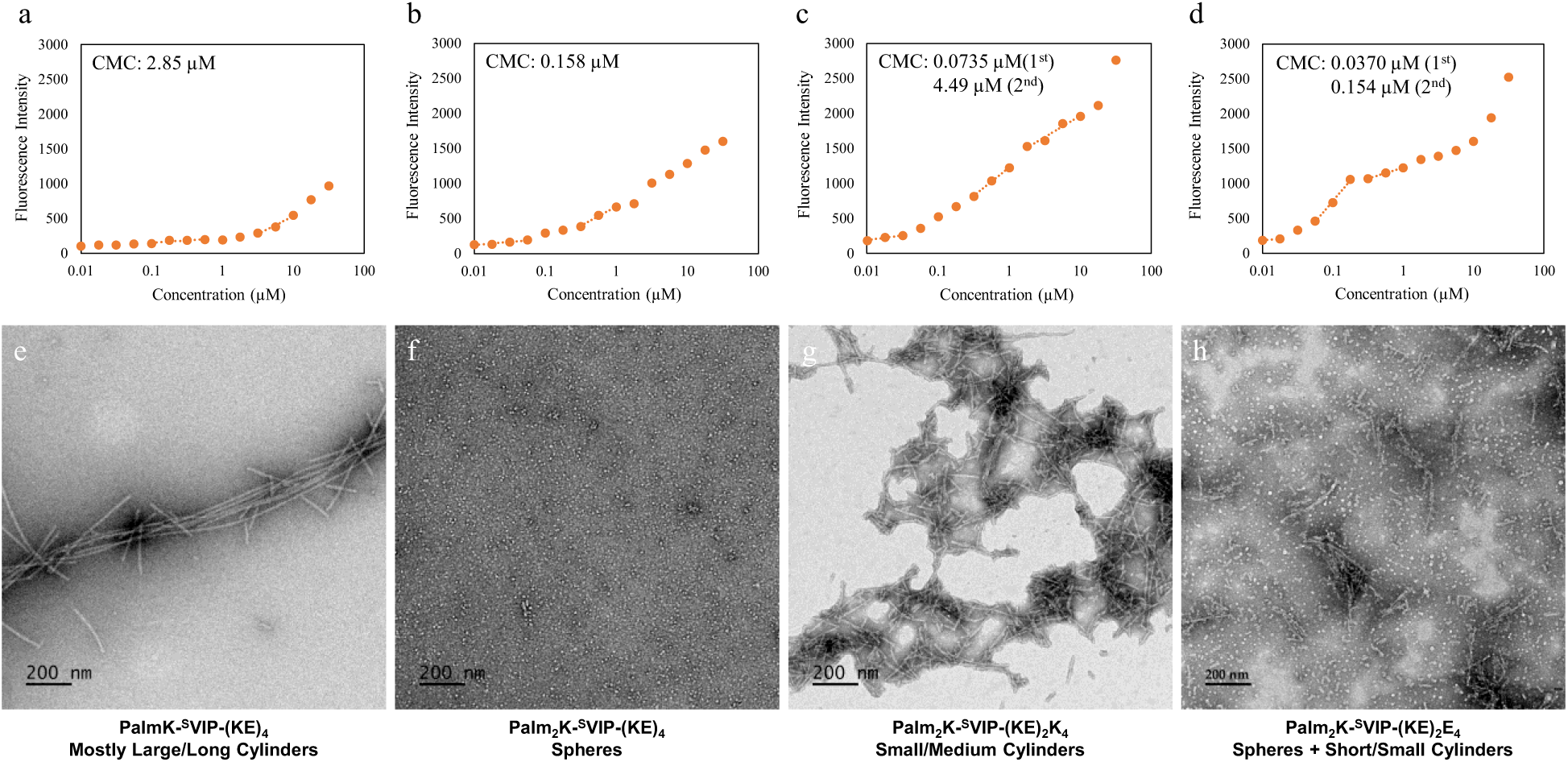
Critical micelle concentration (CMC) and micelle morphologies of PalmK-^S^VIP-(KE)_4_ (a, e), Palm_2_K-^S^VIP-(KE)_4_ (b, f), Palm_2_K-^S^VIP-(KE)_2_K_4_ (c, g) and Palm_2_K-^S^VIP-(KE)_2_E_4_ (d, h). All micrographs were taken at a magnification of 12,000 x with the scale bar for all images representing 200 nm.

**Table 2.** Critical micelle concentration(s) and morphology of VIPAs and ^S^VIPAs.

| VIPAs | CMC | Morphology | <sup>S</sup>VIPAs | CMC | Morphology |
| --- | --- | --- | --- | --- | --- |
| PalmK-VIP-(KE)<sub>4</sub> | 0.333 | Mostly Large/Long Cylinders | PalmK-SVIP-(KE)<sub>4</sub> | 2.85 | Mostly Large/Long Cylinders |
| Palm<sub>2</sub>K-VIP-(KE)<sub>4</sub> | 0.0872 (1<sup>st</sup>)<br>0.409 (2<sup>nd</sup>) | Small/Medium Cylinders | Palm<sub>2</sub>K-SVIP-(KE)<sub>4</sub> | 0.158 | Spheres |
| Palm<sub>2</sub>K-VIP-(KE)<sub>2</sub>K<sub>4</sub> | 0.304 | Mostly Spheres | Palm<sub>2</sub>K-SVIP-(KE)<sub>2</sub>K<sub>4</sub> | 0.0735 (1<sup>st</sup>)<br>4.49 (2<sup>nd</sup>) | Small/Medium Cylinders |
| Palm<sub>2</sub>K-VIP-(KE)<sub>2</sub>E<sub>4</sub> | 0.182 (1<sup>st</sup>)<br>2.72 (2<sup>nd</sup>) | Spheres + Short/Small Cylinders | Palm<sub>2</sub>K-SVIP-(KE)<sub>2</sub>E<sub>4</sub> | 0.0370 (1<sup>st</sup>)<br>0.154 (2<sup>nd</sup>) | Spheres + Short/Small Cylinders |

An additional factor profoundly influencing micelle shape and size is the secondary structure of the peptide^30,31^. Circular dichroism (CD) assessment of VIPAMs and ^S^VIPAMs revealed interesting secondary structure differences between the formulations (**Table 3**). VIP and ^S^VIP were found to contain a near even mix of organized secondary structure (*i.e.*, α-helix and β-sheet - 49.4% and 46.6%, respectively) and disorganized secondary structure (*i.e.*, random coil - 50.6% and 53.4%, respectively) with minimal differences found based on peptide sequence. In contrast to free peptides in solution, all VIPAM and ^S^VIPAM formulations possessed considerably greater organized secondary structure content (*i.e.*, 65.7% - 100%) likely owing to micellization. Confinement of PAs within micelles mimics an artificial tertiary structure which can induce and stabilize organized peptide secondary structure^32^. For single lipid micelles (*i.e.*, PalmK-VIP-(KE)_4_ and PalmK-^S^VIP-(KE)_4_), considerable changes in the distribution of α-helical, β-sheet, and random coil content did little to impact their elongated cylindrical shape. In contrast, double lipid micelles had a clear shape to peptide secondary structure relationship. Double lipid, spherical micelles (*i.e.*, Palm_2_K-VIP-(KE)_2_K_4_ and Palm_2_K-^S^VIP-(KE)_4_) possessed more the greatest combined α-helix and random coil content (*i.e.*, 46.5% and 41.5%, respectively). while double lipid formulations with considerable cylindrical micelle populations (*i.e.*, Palm_2_K-VIP-(KE)_4_, Palm_2_K-VIP-(KE)_2_E_4_, Palm_2_K-^S^VIP-(KE)_2_K_4_, and PalmK-^S^VIP-(KE)_2_E_4_) contained a much lower quantity of these (*i.e.,* 0% - 13.3%). The bulky nature of α-helical and random coil peptide secondary structures has been shown to promote spherical micelle formation, whereas β-sheet structures associated with tighter amphiphile packing and reduced a_o_ values drive cylindrical micelle production^31^. After examining the unique structural relationships between VIPAM and ^S^VIPAM formulations, micelle-cell interactions mediated by VIP/VPAC binding were studied.

**Table 3.** Secondary structure of VIPAMs and ^S^VIPAMs.

| VIPAs | α-Helix | β-Sheet | Random Coil | <sup>S</sup>VIPAs | α-Helix | β-Sheet | Random Coil |
| --- | --- | --- | --- | --- | --- | --- | --- |
| VIP | 8.1% | 41.3% | 50.6% | <sup>S</sup>VIP | 7.9% | 38.6% | 53.4% |
| PalmK-VIP-(KE)<sub>4</sub><br>Mostly Large/Long Cylinders | 18.9% | 81.1% | 0.0% | PalmK-<sup>S</sup>VIP-(KE)<sub>4</sub><br>Mostly Large/Long Cylinders | 33.7% | 32.0% | 34.3% |
| Palm<sub>2</sub>K-VIP-(KE)<sub>4</sub><br>Small/Medium Cylinders | 13.3% | 86.7% | 0.0% | Palm<sub>2</sub>K-<sup>S</sup>VIP-(KE)<sub>4</sub><br>Spheres | 19.7% | 58.5% | 21.8% |
| Palm<sub>2</sub>K-VIP-(KE)<sub>2</sub>K<sub>4</sub><br>Mostly Spheres | 17.3% | 53.8% | 28.9% | Palm<sub>2</sub>K-<sup>S</sup>VIP-(KE)<sub>2</sub>K<sub>4</sub><br>Small/Medium Cylinders | 0.0% | 100.0% | 0.0% |
| Palm<sub>2</sub>K-VIP-(KE)<sub>2</sub>E<sub>4</sub><br>Spheres + Short/Small Cylinders | 12.1% | 87.9% | 0.0% | Palm<sub>2</sub>K-<sup>S</sup>VIP-(KE)<sub>2</sub>E<sub>4</sub><br>Spheres + Short/Small Cylinders | 0.0% | 100.0% | 0.0% |

### Cell Association and Uptake of Micelles with Varying Chemical and Physical Properties

To investigate the influence VIPAMs and ^S^VIPAMs with different shapes and sizes have on Mφs, initial (*i.e.*, 1 hour) and prolonged (*i.e.*, 6 hours) micelle/cell co-incubation was observed using confocal microscopy (**Figure 6** and **Figure 7**). Green dot clusters located between the cell nucleus (blue) and cell membrane (red) were indicative of micelle internalization and denoted with yellow arrows. In contrast, yellow-green dot masses found co-localized with the cell membrane were considered surface-associated and highlighted with grey arrows. At the earlier time point, VIPAM chemistries capable of forming spherical micelles (*i.e.*, Palm_2_K-VIP-(KE)_2_K_4_ and Palm_2_K-VIP-(KE)_2_E_4_) were found to be both associated with the surface of Mφs as well as between the membrane and nucleus (**Figure 6c-d**). Conversely, VIPAMs which formed cylindrical micelles (*i.e.*, Palm_2_K-VIP-(KE)_4_ and PalmK-VIP-(KE)_4_) were only found along the Mφ cell membrane (**Figure 6a-b**). At 6 hours, Mφ-internalized micelles were found for all VIPAM chemistries that generated spherical and more compact cylindrical micelles (**Figure 6f-h**) whereas the elongated cylindrical micelles made by PalmK-VIP-(KE)_4_ remained associated solely with the cell surface (**Figure 6e**). Mφ uptake of ^S^VIPAM formulations that formed spherical and more compact cylindrical micelles (*i.e.*, Palm_2_K-^S^VIP-(KE)_4_, Palm_2_K-^S^VIP-(KE)_2_E_4_, and Palm_2_K-^S^VIP- (KE)_2_K_4_) was observed at both time points (**Figure 7b-d, f-h**). In contrast, limited Mφ association with and no internalization of elongated cylindrical PalmK-^S^VIP-(KE)_4_ micelles was observed throughout the experiment (**Figure 7a,e**). The rapid internalization of spherical and short cylindrical micelles independent of peptide chemistry (**Figure 6c-d** and **Figure 7c-d**) aligns with effects expected from endocytosis of extracellular material under 100 nm in size^33^. Specifically, multiple endocytic pathways capable of internalizing extracellular material within this size range lead to no advantage of VPAC targeting by VIPAMs over ^S^VIPAMs. Interestingly, small/medium cylindrical micelle-inducing formulations possessed chemistry-dependent internalization speed with VIPAMs being uptaken by Mφs more slowly than ^S^VIPAMs (**Figure 6b,f** and **Figure 7c,g**). Previous research has shown that intermediate-sized particles (*i.e.*, 300 - 3000 nm) are capable of fitting into the ruffles of the APC cell membrane more easily than their smaller or larger counterparts^12,19,34,35^ and can be readily internalized by phagocytosis^13,35^. While ^S^VIPAMs would be able to follow this relatively quick uptake pathway, VIPAMs likely bind their cognate surface receptor (*i.e.*, VPAC) localizing them there for longer periods of time. Previous research has found that Mφs can internalize materials by phagocytosis within an hour^36^, whereas significant VPAC internalization due to VIP binding^37^ may take longer^38–40^ and be inefficient for the internalization of intermediate-sized particles^41^. The chemistry-dependent association of long cylindrical micelles (> 3000 nm) is unsurprising as these would be too large to fit into the cell membrane ruffles, so only those that could directly bind the cell surface (*i.e.*, VIPAMs to VPAC) would be capable of co-localizing with cells (**Figure 6a,e**). Conversely, those that could not bind VPAC (*i.e.*, ^S^VIPAMs) would be unable to facilitate prolonged contact with cells (**Figure 7a,e**). The lack of internalization by surface-associated long cylindrical VIPAMs is expected as their size would require considerable cell membrane movement and reorganization which is a time- and labor-intensive process^13^. To better quantify the interesting effects that micelle shape and size have on cell association, the interactions VIPAMs and ^S^VIPAMs have with co-incubated Mφs were also analyzed by flow cytometry.

**Figure 6.**
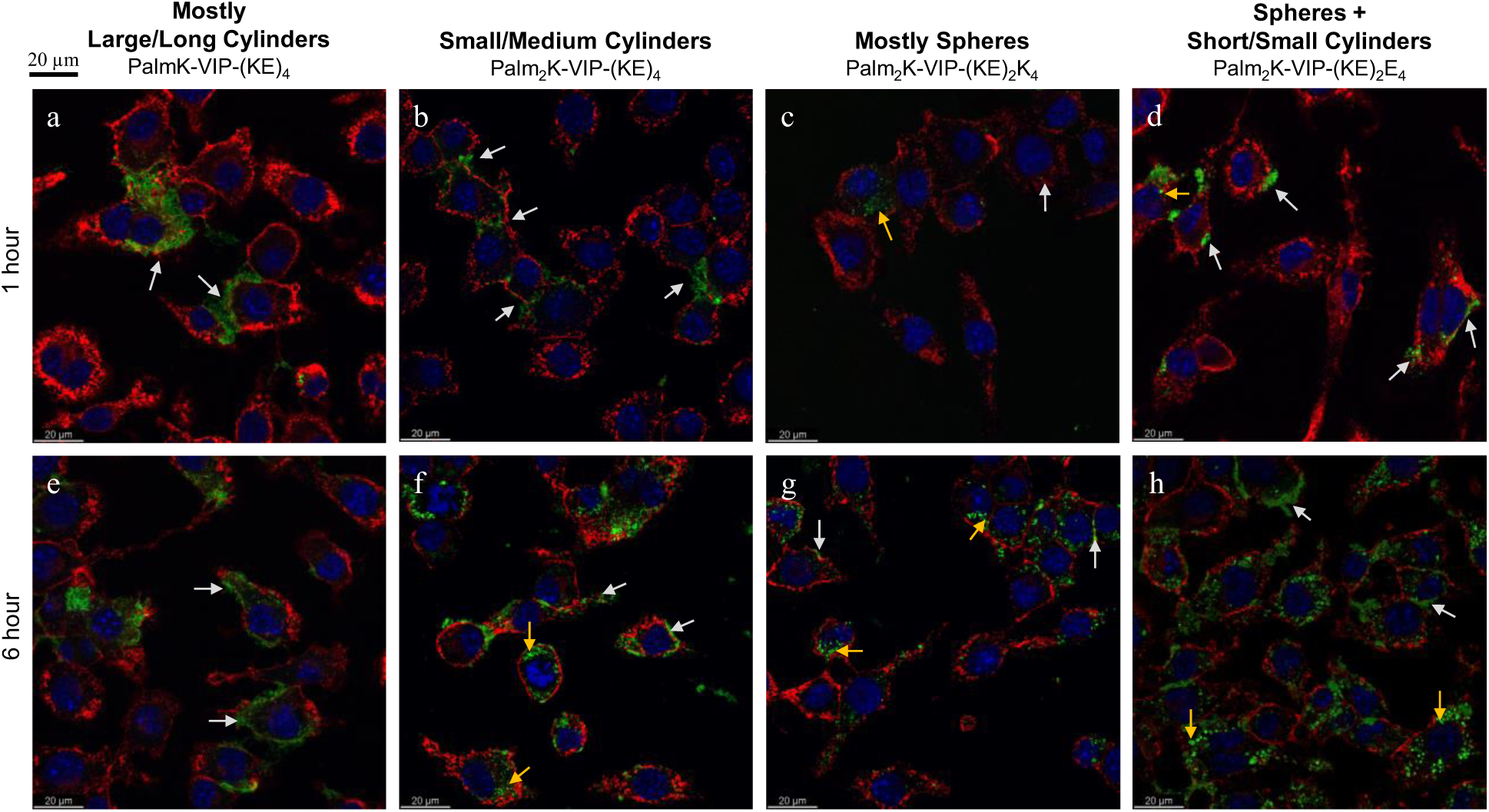
Representative confocal microscopy images of VIPAM association with Mφs at 1 hour (a-d) and 6 hours (e-h). The cell membrane and cell nucleus were stained by WGA (red) and DAPI (blue), respectively, whereas the FAM-labeled micelles appear in green. Yellow arrows indicate internalized micelles and grey arrows indicate micelles associated with the cell surface. Images were taken using a 40 X lens for which the scale bar is equal to 20 µm.

**Figure 7.**
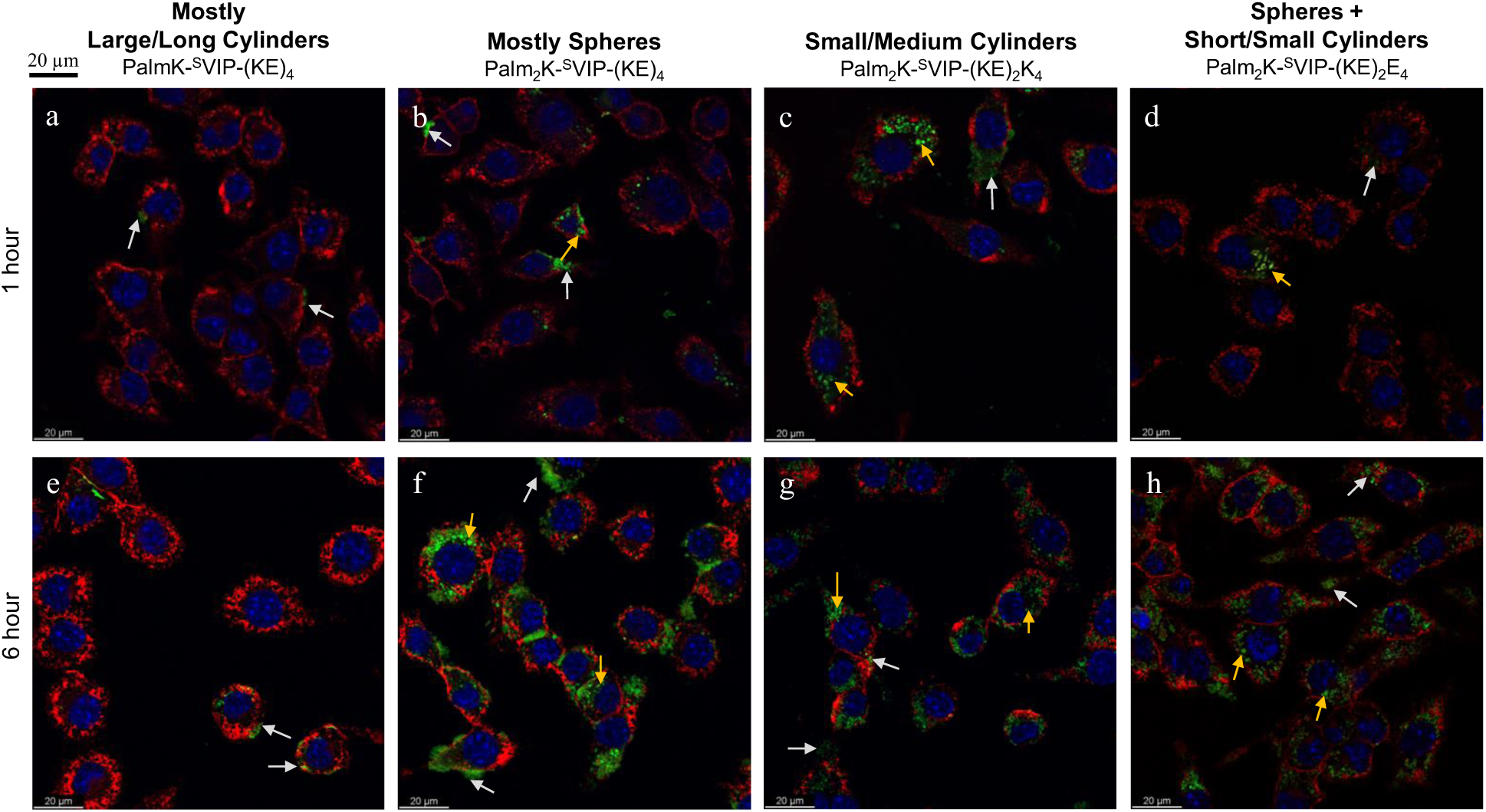
Representative confocal microscopy images of ^S^VIPAM association with Mφs at 1 hour (a-d) and 6 hours (e-h). The cell membrane and cell nucleus were stained by WGA (red) and DAPI (blue), respectively, whereas the FAM-labeled micelles appear in green. Yellow arrows indicate internalized micelles and grey arrows indicate micelles associated with the cell surface. Images were taken using a 40 X lens for which the scale bar is equal to 20 µm.

Mφs were co-cultured with all micelle formulations for one and six hours, and the percentage of Mφs associated with fluorophore-labeled micelles as well as their median fluorescence intensity (MFI) were measured (**Figure 8-9**). Unsurprisingly, micelle peptide specificity was found to play a crucial role in facilitating rapid cell association as Mφs were 2 - 10 times more likely to be associated with VIPAMs than their analogous ^S^VIPAMs at the early time point (**Figure 8a**). In addition, all tested VIPAMs, though possessing different morphologies, facilitated a similar percentage of micelle-associated Mφs suggesting initial association was mainly regulated by cell surface receptor specificity regardless of micelle shape and size. Without this receptor-mediated effect, Mφs preferred to associate with small/medium cylindrical ^S^VIPAMs (*i.e.*, Palm_2_K-^S^VIP-(KE)_2_K_4_) which can be efficiently entrapped in Mφ membrane ruffles. The MFI of fluorophore-labeled micelle-associated Mφs was relatively low for all formulations after 1 hour of co-incubation suggesting only a few micelles were associated with Mφs at this early time point (**Figure 8b**). Large/long cylindrical VIPAMs displayed a slightly higher MFI relative to all other formulations resulting from their increased fluorescent signal per micelle despite low surface attachment.

**Figure 8.**
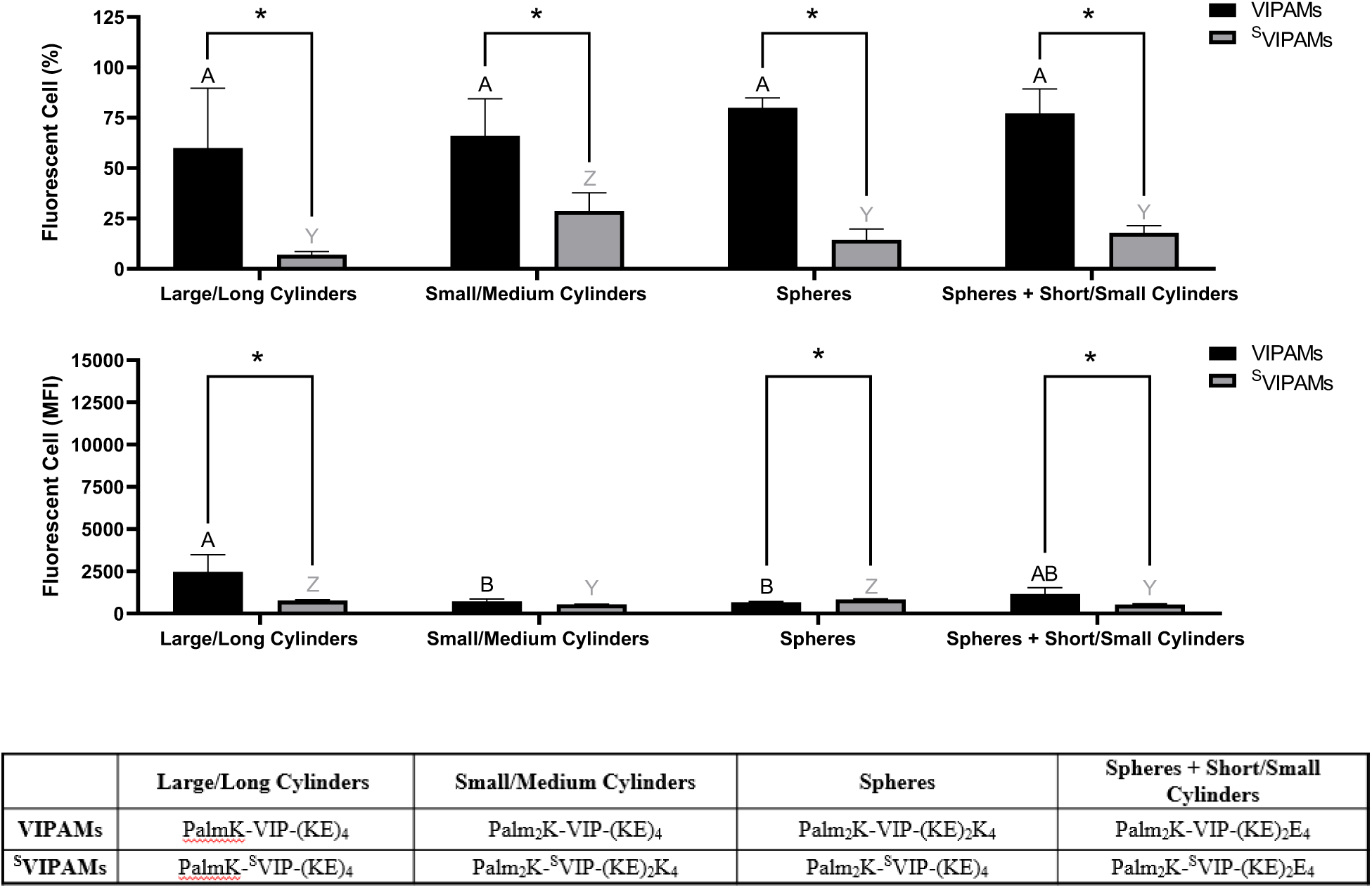
Fluorescent cell population related to total cell population (a) and mean fluorescence intensity (MFI) (b) of Mφs incubated for 1 hour with VIPAMs or ^S^VIPAMs of different shapes and sizes. In each graph, groups possessing the same letter or not possessing a star between them have no statistically significant difference (p > 0.05). Black (A - B) and grey (Y - Z) letters were used respectively to compare VIPAMs and ^S^VIPAMs with different shapes and sizes, while * was used to compare VIPAMs and ^S^VIPAMs that possessed the same shape and size.

**Figure 9.**
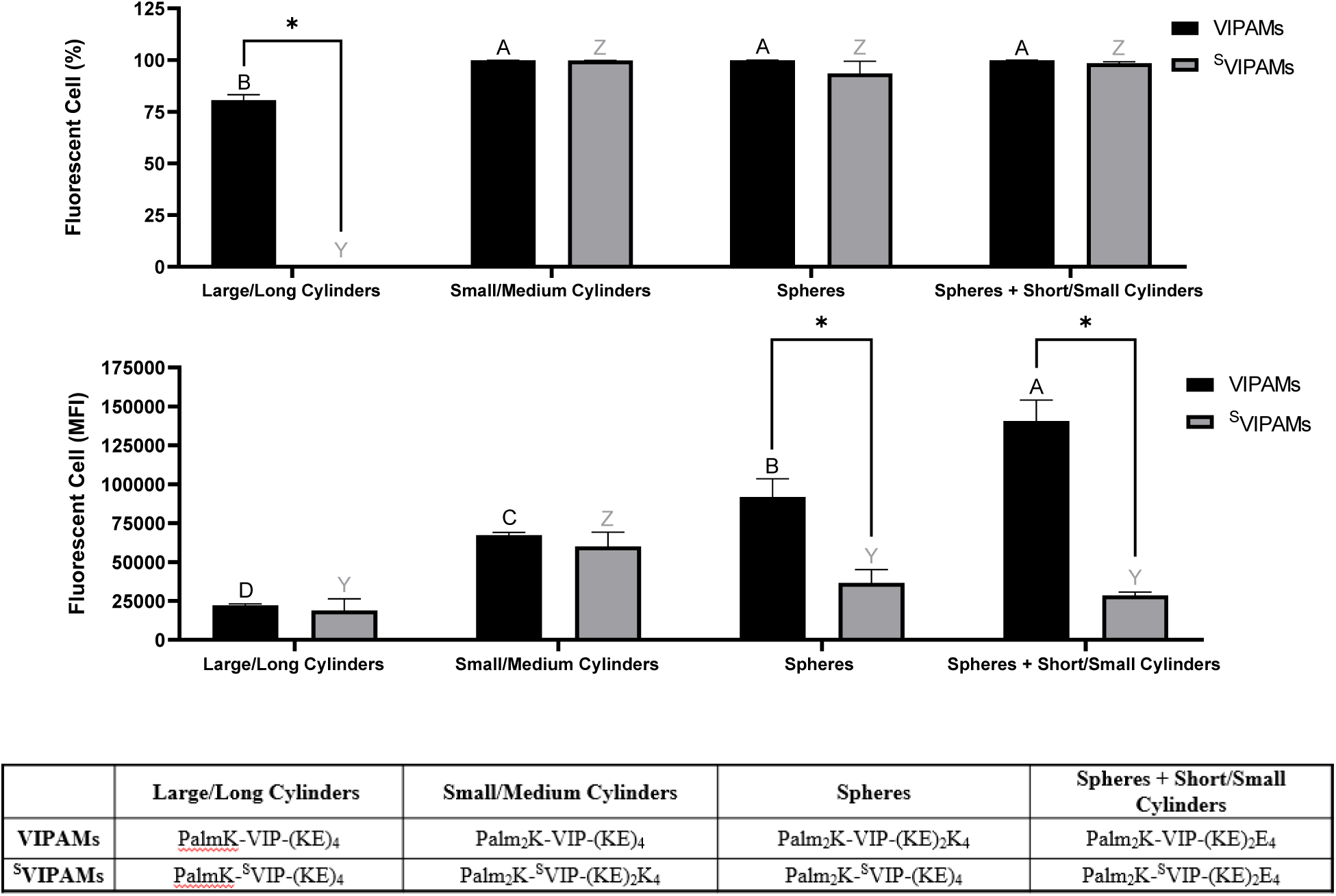
Fluorescent cell population related to total cell population (a) and mean fluorescence intensity (MFI) (b) of Mφs incubated for 6 hours with VIPAMs or ^S^VIPAMs of different shapes and sizes. In each graph, groups possessing the same letter or not possessing a star between them have no statistically significant difference (p > 0.05). Black (A - B) and grey (Y - Z) letters were used respectively to compare VIPAMs and ^S^VIPAMs with different shapes and sizes, while * was used to compare VIPAMs and ^S^VIPAMs that possessed the same shape and size.

At 6 hours, almost all Mφs were associated with co-cultured micelles regardless of their chemistry, shape, or size, besides large/long cylindrical ^S^VIPAMs (*i.e.*, PalmK-^S^VIP-(KE)_4_) (**Figure 9a**). Nearly 75% of Mφs were found associated with long cylindrical VIPAMs, but their low MFI suggests low numbers of micelles were present per cell with each micelle contributing a large fluorescent output. The large size and non-specificity of long cylindrical ^S^VIPAMs greatly limited their association with Mφs and engulfment by Mφ membrane ruffles^13,19^ as well, aligning with prior data (**Figure 7e**). The quantity of micelle association with Mφs, as reported by MFI, showed that VIPAM formulations generating solely spherical (*i.e.,* P_2_K-VIP-(KE)_2_K_4_) or solely small/medium cylindrical micelles (*i.e.,* Palm_2_K-VIP-(KE)_4_) had significantly higher MFIs compared with long cylinders, coinciding with their smaller size and improved ability to associate with and cross the cell membrane (**Figure 9b**). Furthermore, spherical micelles displayed a greater MFI over small/medium cylindrical micelles. Although less is known about the size of VPAC receptors, typical GPCr transmembrane segments have diameters capable of accommodating very small micelles^42^. Because spherical micelles are made of less individual PAs, a greater number of total micelles would be formed upon reconstitution. Spherical micelles might then maximize VPAC receptor occupation relative to other VIPAM formulations as more micelles capable of interacting with their cognate receptors would be expected at any single time point. Interestingly, the greatest Mφ association was found with Palm_2_K-VIP-(KE)_2_E_4_ which forms a mix of spherical and short/small cylindrical micelles, potentially reflecting the impact of PAM size on Mφ association as well as the importance of fluorophore-labelled PA content per micelle. In contrast to spherical micelles, the largest-sized VIPAMs (*i.e.*, Palm_2_K-VIP-(KE)_4_ and PalmK-VIP-(KE)_4_) contain much more PAs per micelle and thus possess more fluorophore-labelled PAs per micelle (∼ 15 - 1,500) as well, compared to the ∼ 2 - 4 expected for spherical micelles. Association events between these bigger VIPAMs and VPAC receptor(s), though less in frequency, would yield a much higher fluorescent payload, though large micelle size might reduce Mφ VPAC occupancy due to shielding of unbound receptors. Palm_2_K-VIP-(KE)_2_E_4_ forms both spherical and small cylindrical micelles, which may function synergistically to increase the number of receptor binding events and total delivered fluorescent payload. The sizeable spherical PAM population can accommodate many cell surface receptors, while the cylinders observed in this formulation may possess a somewhat high fluorophore per micelle content (∼ 3 - 50), enhancing the resulting fluorescent signal. Moreover, their smaller size might circumvent the problem of shielding unbound receptors leading to efficient VPAC binding. This data corresponds well to the confocal microscopy results which showed co-incubated Mφs had their cell membranes coated by Palm_2_K- VIP-(KE)_2_E_4_ micelles (**Figure 6h**) to a much greater extent than all other VIPAM formulations (**Figure 6e-g**).

Without cell surface receptor specificity, the MFI of Mφs associated with ^S^VIPAMs remained lower than or similar to that of Mφs contacting VIPAMs. All ^S^VIPAM formulations except one demonstrated insignificantly different levels of cell interactions, with small/medium cylindrical ^S^VIPAMs displaying an enhanced level of fluorescence similar to that of the structurally analogous VIPAMs. The relatively high uptake of nonspecific small/medium cylinders aligns with previous observations (**Figure 7f**). With this formulation, VPAC specificity may slow down PA and PAM internalization over nonspecific micelles by localizing VIPAMs at receptors^38–40^ preventing efficient cellular entry by phagocytosis. Comparing cell association across micelles with different peptide chemistry showed that all VIPAMs yielded greater or similar MFI values relative to their ^S^VIPAM counterparts indicating the overall value of VPAC-based Mφ targeting.

## Conclusion

This research provides significant insight into how modifications in lipid content and peptide sequence can be leveraged to generate PAMs with varying shapes and sizes which directly influences their association to and uptake by Mφs. Micelle morphology was greatly influenced by lipid tail steric hindrance as well as more finely manipulatable by minor changes in hydrophilic peptide block amino acid content. Interestingly, altering the position of the amino acids within the bioactive peptide region (*i.e.*, using ^S^VIP compared to VIP) modulated which double lipid PAs formed various nanostructures though batches of similar micelle architectures were able to be made regardless of whether VIP or ^S^VIP was incorporated. Co-incubation of these various micelle formulations with Mφs revealed that presentation of cell targeting VIP over its scrambled peptide analog (*i.e.*, ^S^VIP) dictated rapid association with VIP, more so than micelle shape and size. That being said, micelle prolonged association with and internalization by Mφs was likely influenced by a combination of biomolecular material structure and chemistry as well as route of entry into cells. Although all VIPAMs demonstrated greater or similar levels of cellular association compared to ^S^VIPAMs, structurally dissimilar micelles interacted with cells at different rates indicating potential differences in internalization mechanisms. Smaller VPAC-specific micelles present in larger numbers may bind a large quantity of VPAC receptors while also exploiting other mechanisms of nonspecific internalization^37–40^. By contrast, slightly larger small/medium cylindrical micelles not specific to VPAC may actually benefit from non-specificity as these are more rapidly internalized by phagocytosis than VPAC-specific, smaller or larger particles^12,13,19,33–35^. From these studies, Palm_2_K-VIP-(KE)_2_E_4_ was found to produce a mix of spherical and short/small cylindrical micelles from 10 - 300 nm in largest dimension, which achieved the best Mφ association being able to be both internalized and persist on the cell surface. While exciting, the influence micelle receptor specificity and material properties have on their bioactivity still needs to be further studied to help generate more detailed chemistry-structure-function relationships for immunomodulatory materials as well as specifically down select the formulation best capable of inducing productive and sustained anti-inflammatory effects for future clinical applications.

## Notes

### Competing Interest Statement

The authors have declared no competing interest.

